# The 4T1 breast carcinoma model possesses hybrid epithelial/mesenchymal traits linked to a highly metastatic phenotype

**DOI:** 10.1101/2024.02.02.578710

**Authors:** Mary E. Herndon, Katherine Gibson-Corley, Lori L. Wallrath, Michael D. Henry, Christopher S. Stipp

**Author notes:** Deceased (Mary E. Herndon).

## Abstract

Epithelial-mesenchymal transitions (EMTs) have emerged as a mechanism for carcinomas to gain metastatic capabilities. As classically defined, EMTs entail downregulation of E-cadherin, viewed as a master enforcer of an epithelial phenotype, and upregulation of mesenchymal markers such as N-cadherin and vimentin. Contrary to this, E-cadherin appears to be retained in many invasive carcinomas and promote collective cell invasion. Therefore, major questions remain concerning the role of E-cadherin in metastasis. To investigate how E-cadherin regulates metastasis, we selected murine 4T1 breast carcinoma cells, a widely utilized model of stage IV metastatic breast cancer that retains E-cadherin expression. Using RNA interference and constitutive expression, we demonstrate that the expression level of E-cadherin does not determine 4T1 metastatic capacity in mice. Despite high levels of expression, E-cadherin is unable to confer an epithelial phenotype with stable organized cell-cell junctions. Moreover, orthotopic 4T1 tumors in mice display co-expression of E-cadherin and vimentin and contain subregions of EMT-like loss of E-cadherin. In addition, we find that 4T1 cells co-express epithelial and mesenchymal isoforms of p120-catenin. These findings support 4T1 cells as a model for carcinomas that possess hybrid traits of epithelial and mesenchymal states that promote invasion and metastasis. As such, the 4T1 model provides a platform for investigating strategies to reinstate E-cadherin’s ability to promote stable cell-cell junctions in E-cadherin-positive cancers, and for identifying which aspects of an epithelial phenotype may serve to facilitate the macroscopic growth of metastatic colonies.

## Introduction

Epithelial-mesenchymal transitions (EMTs) have become a framework for explaining the metastatic capacity of carcinomas (tumors that arise in epithelial cell layers) [1]. As extensively documented by foundational studies, EMT programs, which also play critical roles in development, wound healing, and fibrosis, are governed by EMT transcription factors, including Zeb1, Zeb2, Twist, and Snail [2–5]. As originally described, features of mesenchymal cancer cells produced by aberrant EMT activation in tumor progression included (i) loss of the epithelial cell-cell adhesion protein, E-cadherin, (ii) upregulation of mesenchymal cadherins, such as N-cadherin, (iii) loss of epithelial cytokeratin intermediate filaments, (iv) upregulation of the mesenchymal intermediate filament, vimentin, (v) disrupted cell-cell cohesion and loss of apical-basal polarity, and (vi) increased cell migration and invasion into connective tissue, resulting in tumor cell dissociation from the primary tumor and dissemination to distant locations [2–5]. Subsequently, intriguing connections between EMT events and the acquisition of stem cell-like phenotypes were identified, with important implications for chemoresistance, recurrence, and the capacity of disseminated tumor cells to generate macroscopic metastatic tumors [6–10]. In addition, EMT-like states may also contribute to intermediate steps in the metastatic process, such as trans-endothelial migration of circulating tumor cells [11].

Early debates on the role of EMTs in metastasis often centered around the apparent lack of cancer cells with a definitive mesenchymal phenotype in human clinical carcinoma specimens [12]. A hypothesis for the seeming paucity of mesenchymal cancer cells at either primary or metastatic tumor sites is that EMTs occur in a small subset of primary tumor cells and that a reverse, mesenchymal-to-epithelial transition (MET) occurred during outgrowth of macroscopic metastatic tumors [13]. This hypothesis has garnered considerable experimental support [14–21]. With the advent of the ability to capture circulating tumor cells (CTCs) and to profile gene expression via single cell RNA seq, another hypothesis for the elusiveness of mesenchymal carcinoma cells is that a hybrid epithelial/mesenchymal (E/M) phenotype, also called quasi-mesenchymal phenotype, is linked to the acquisition of metastatic capacity more than a purely mesenchymal phenotype [9,22].

In many respects, debates about the functional role of E-cadherin in regulating metastasis have mirrored the debates about the roles of EMT and MET. On one hand, E-cadherin has been described as “the keystone of the epithelial state” [23], such that loss of E-cadherin on its own is sufficient to trigger an EMT phenotype [24], while enforced E-cadherin expression may be sufficient to prevent metastatic outgrowth [25,26]. On the other hand, E-cadherin can play a key role in collective cell migration and invasion [27], and may function in some settings to promote the metastatic outgrowth of tumor cells *in vivo* [28].

The immune-competent, murine 4T1 cell model of stage IV breast cancer has been an embodiment of both the progress and the lively debates in the EMT field. For example, in one early study, highly metastatic 4T1 cells provided an example of the important role of the EMT master regulator, Twist, in driving metastasis [29]. However, subsequent studies have noted E-cadherin expression by 4T1 cells, despite their highly metastatic phenotype, and suggested possible pro-metastatic roles for E-cadherin in the 4T1 model system [30,31].

In this study, we specifically sought to clarify the role of E-cadherin in the metastatic capacity of the widely used 4T1 model. Surprisingly, we find that downregulation of E-cadherin, sufficient to disrupt cell-cell cohesion, had minimal impact (positive or negative) on metastatic capacity. Further, we find that despite the presence of E-cadherin, 4T1 epithelial cell-cell junctions are dynamic and unstable, leading to highly migratory phenotypes in 2D and 3D models of tumor cell migration and invasion. In an orthotopic *in vivo* mouse model, we find evidence of overt EMT-like events of 4T1 cell metastasis. Taken together, these findings are consistent with the highly aggressive phenotype in this widely utilized model of breast cancer progression. Moreover, our findings help to explain the mechanisms by which 4T1 cells have the capacity to manifest an aggressive hybrid E/M phenotype associated with metastasis.

## Materials and Methods

### Antibodies

Primary antibodies used in the study are listed in Supplementary Table 1. Fluorescently labeled secondary antibodies for immunostaining and western analyses were from Thermo Fisher Scientific (Waltham MA, Alexa Fluor dyes) and Rockland Immunochemicals, Inc, (Pottstown, PA, IR dyes).

### Cell culture, RNA interference, and retroviral transduction

All cell culture reagents and antibiotics were from Thermo Fisher Scientific (Waltham, MA), except where otherwise specified. The 4T1 cell line was obtained from the American Type Culture Collection (ATCC; Manassas, VA), and the 4T07, 67NR, and 168FARN mouse mammary carcinoma lines were gifts from Fred Miller (Wayne State University). Collectively, these cells were cultured in RPMI supplemented with 1X NEAA. A431 epithelial carcinoma cells (ATCC; Manassas, VA) and GP2-293 cells (Takara Bio USA; San Jose, CA) were cultured in high glucose DME. All cultures were supplemented with 2 mM L-glutamine, 100 U/ml penicillin, 100 μg/ml streptomycin, and 10% fetal bovine serum (Valley Biomedical Inc.; Winchester, VA). Serum free medium (SFM) for the mammary carcinoma lines was RPMI supplemented with 1X NEAA, 25 mM HEPES and 5 mg/ml cell culture grade BSA (Sigma-Aldrich, product number A1470; St. Louis, MO). Antibiotics used for selection included G418, zeocin, and puromycin (Gold Biotechnology; St. Louis, MO).

To facilitate monitoring tumor growth *in vivo*, 4T1 cells were transduced with a *luciferase* cDNA cloned into the pQCXIN retroviral expression vector (Takara Bio USA; San Jose, CA), selected with 0.5 mg/ml G418, and maintained in 0.1 mg/ml G418. This parental cell line (4T1-lucIN) was used to create E-cadherin knockdown cells (Ecad-KD), cells constitutively expressing myc-tagged E-cadherin (Ecad-myc), and empty vector control (VEC) cells.

For RNAi, double-stranded oligonucleotides encoding short hairpin RNAs (shRNAs) targeting the mouse *E-cadherin* mRNA were annealed and then cloned into the pSIREN-RetroQ retroviral vector (BD Biosciences; Franklin Lakes, NJ). The shRNA targeting sequence for *E-cadherin* was 5’-TACATCCTTCATGTGAGAGTG-3’. This construct was co-transfected with the pVSV-G retroviral coat protein expression vector into GP2-293 packaging cells using Effectene (Qiagen Sciences, Inc.; Germantown, MD). At 24 h and again at 48 h post-transfection, virus-containing medium was collected, 0.45 µ filtered, supplemented with 4 μg/ml polybrene (Sigma-Aldrich, St. Louis, MO), and then used to transduce 4T1-lucIN cells. Stable transductants (Ecad-KD cells) were selected with 2 µg/ml puromycin, maintained in 1 µg/ml puromycin and 0.1 mg/ml G418, and sorted by flow cytometry to obtain a polyclonal population with greatly reduced E-cadherin expression.

To create Ecad-myc cells, mouse E-cadherin cDNA (Image Clone #3002385; Open Biosystems, Inc., which was acquired by Thermo Fisher Scientific, Waltham, MA), was cloned into an LXIZ retroviral vector engineered to contain a high affinity 3’-Myc tag [32]. The same LXIZ vector lacking an insert was used to create empty vector control cells (VEC). These constructs were transfected into GP2-293 cells, then used to transduce 4T1-lucIN cells by the method above. Stable transductants were selected with 0.3 mg/ml zeocin, and then maintained in 0.1 mg/ml zeocin and 0.1 mg/ml G418.

### Immunoprecipitation and western analysis

Cells were lysed in PBS with 1% detergent, protease inhibitors (2 mM PMSF, 10 µg/ml aprotinin, 5 µg/ml leupeptin and 5 µg/ml E-64) and HALT phosphatase inhibitor (Thermo Fisher Scientific, Waltham, MA). Detergents were NP-40 or Brij 96V (both from Sigma-Aldrich, St. Louis, MO). In some experiments, cells were cell-surface biotinylated with 0.1 mg/ml Sulfo-NHS-LC Biotin (Thermo Fisher Scientific, Waltham, MA) in HBSM (20 mM HEPES pH 7.2, 150 mM NaCl, 5 mM MgCl_2_) for 1 hour at room temperature and then rinsed three times with HBSM prior to lysis. Lysates were clarified, protein concentrations measured using the Red 660 Protein Assay (G-Biosciences; St. Louis, MO), then lysate concentrations were normalized. Immune complexes, composed of primary antibodies combined with Protein G Agarose (Thermo Fisher Scientific, Waltham, MA) were collected, separated by SDS-PAGE, and transferred to nitrocellulose. The resulting membranes were blocked with Intercept TBS (LI-COR Biosciences, Lincoln, NE), and incubated with primary antibodies diluted in 10% Intercept TBS diluted in TBST (20 mM Tris pH 7.5, 150 mM NaCl, 0.1% Tween-20). The membranes were incubated with appropriate fluorescently labeled secondary antibodies diluted in 10% Intercept TBS/TBST and analyzed with a LI-COR near-infrared fluorescence imager. Biotinylated proteins were detected with NeutrAvidin DyLight 800 (Thermo Fisher Scientific, Waltham, MA).

### Immunostaining cultured cells and spheroids

4T1 and related cell lines were plated on sterile uncoated, rat tail collagen I (Thermo Fisher Scientific, Waltham, MA) coated, or A431 cell-matrix conditioned glass cover slips. For A431 conditioned cover slips, A431 cells were plated at confluence on glass cover slips in 24 well plates, allowed to grow overnight, and removed by treating with PBS/1 mM EDTA at 37° for 30 min followed by several washes with PBS/EDTA until cover slips were cleared. Cells cultured on cover slips were fixed with 10% formalin in HEPES-buffered saline (HBS) with 4% sucrose and 1 mM MgCl_2_, rinsed twice with TBS, and blocked with 10% goat serum in PBS. For staining intracellular epitopes, cells were permeabilized with the addition of 0.1% NP-40 during the blocking step. Cells were stained for 1 hour with primary antibodies in blocking buffer, briefly washed four times with TBST, followed a 45-minute treatment with the appropriate fluorescently labeled secondary antibodies in blocking buffer. For F-actin staining, Alexa Fluor 594 phalloidin (Thermo Fisher Scientific, Waltham, MA) was included with the secondary antibody in TBST/0.1% NP-40 instead of blocking buffer. For nuclear staining, 0.5 µg/ml DAPI (Sigma-Aldrich, St. Louis, MO) was included in the secondary antibody step. After four additional washes with TBST, cover slips were mounted with Prolong Gold (Thermo Fisher Scientific, Waltham, MA) and analyzed by fluorescence microscopy. For immunostaining of tumor spheroids, spheroids created as described below were embedded in 0.8 mg/ml rat tail collagen I, in a glass bottomed 35 mm dish. After 48 hours of invasion, spheroids were fixed for two hours at room temperature with 10% formalin in PBS, rinsed extensively with TBS, and blocked overnight in PBS with 10% goat serum.

Primary antibody was added in block and incubated overnight at 4°C. Spheroids were rinsed extensively with PBS, with the last rinse overnight at 4°C. Secondary antibody was added in block and incubated overnight at 4°C, followed by rinsing as for the primary antibody.

Following the last rinse, spheroids were post-fixed for two hours at room temperature with 10% formalin and then rinsed with TBS. Fluorescent images were acquired through the glass bottomed dish using a 20X objective.

### Tumor spheroid assays

To create spheroids for 3D growth and invasion assays, 1 x 10^4^ cells per spheroid were plated in 96 well V-bottom plates that had been coated twice by air-drying 25 µl of 20 mg/ml polyhydroxyethylmethacrylate (poly-HEMA; Sigma-Aldrich, St. Louis, MO) in 95% ethanol in each well. Plates were spun briefly in a tabletop centrifuge to settle the cells, then cultured overnight in standard growth medium to allow spheroid formation. To create collagen gels for the invasion assay, 700 µl per well of 0.8 mg/ml rat tail collagen I in RPMI was allowed to polymerize for 1 h at 37°C in a 12 well plate. The plate was then placed on ice, and the spaces between the wells were filled with ice cold PBS to maintain temperature during the transfer of spheroids. A second 700 µl of ice cold 0.8 mg/ml collagen I was overlaid on the cushion of pre-polymerized collagen I, and 6 spheroids per cell type were transferred to each collagen-containing well. To transfer, spheroids were recovered from the V-bottom wells by gently pipetting with a yellow tip to dislodge and capture the spheroids, and then spheroids were allowed to settle by gravity out of the tip and into the collagen, without ejecting any culture medium. After transfer, the collagen was polymerized for one hour at 37°C, and wells were overlaid with 1 ml of SFM. Spheroids were photographed with a 4X objective at indicated time points. To compare spheroid integrity of parental, E-cadherin forced expression, and E-cadherin-silenced cell lines, spheroids formed as above were recovered by gentle pipetting and plated in 35 mm dishes for immediate photography.

### Time-lapse video-microscopy

For random cell migration in 2D, a total of 1 X 10^5^ 4T1 parental cells were plated in standard growth medium on 35 mm culture dishes containing A431-conditioned substrate prepared as described above. The next day, the culture was maintained on a Leica DMIRE2 inverted microscope in a stage incubator (20/20 Technology, Inc., Wilmington, NC) providing a humidified 5% CO_2_, 37°C atmosphere. OpenLab software running on an Apple iMac computer-controlled illumination and image acquisition. Images were acquired at a rate of one frame every three min for six hours, using a Hamamatsu ORCA-285 CCD camera and a 20X C Plan phase objective.

For tumor spheroid invasion in 3D, spheroids created as described above were embedded in 0.8 mg/ml rat tail collagen I in glass bottomed 35 mm dishes, instead of 12 well plates. Cell invasion was monitored with a 4X objective at a rate of one frame every five minutes for 40 hours.

### Breast cancer growth and metastasis *in vivo*

All animal procedures in this study were approved by the University of Iowa Animal Care and Use Committee, Iowa City, IA (Approval No. 5031328). Isoflurane was used for procedures requiring anesthesia, and euthanasia was performed by CO_2_ inhalation followed by cervical dislocation. Female BALB/c mice (NCI-Frederick) were implanted with 5,000 cells in a volume of 50 μL in the fourth mammary fat pad. Bioluminescent imaging (BLI) was conducted in an IVIS100 imaging system (Perkin Elmer, Shelton, CT) after intraperitoneal injection of luciferin (100 μL of 15 mg/mL solution/10 g mouse body weight) as described previously. Whole body tumor growth rates were measured as follows: a rectangular region of interest was placed around the dorsal and ventral images of each mouse, and total photon flux (photons/sec) was quantified using Living Image software v2.50. The dorsal and ventral values were summed and mean BLI values for each group were plotted weekly. Primary tumor growth was also measured by caliper, and tumor volumes were calculated using the formula ½(L x W^2^). For semi-quantitative analysis of spontaneous metastasis to lung, *ex vivo* BLI was conducted on lungs harvested at assay endpoint (day 28). To ensure the best possible uniformity of measurement conditions, mice were sacrificed in groups of 2 or 3, and lungs were immediately harvested and imaged *ex vivo*. All lungs were imaged within 20 to 30 minutes after euthanasia. 4T1 cells colonizing the lungs were recovered by mincing the lungs with a sterile razor blade and digesting with 200 U/mL collagenase II (Worthington Biochemical Corp.; Lakewood, NJ) in complete medium for 15 minutes at 37°C. Explanted cells were grown out under G418 selection for analysis of E-cadherin expression by cell surface labeling and immunoprecipitation, as described above. Some lungs were harvested at the endpoint and fixed in 4% paraformaldehyde overnight at 4°C, rinsed and transferred to 30% ethanol, and stored at 4°C for later histological analysis.

### Histological analysis

Formalin fixed paraffin embedded tissues were routinely stained with hematoxylin-eosin (H&E) (Sigma-Aldrich, St. Louis, MO), and immunohistochemistry was performed for E-cadherin and vimentin. Antigen unmasking of paraffin sections was performed (citrate buffer, pH 6) in a de-cloaker. Endogenous peroxidase activity was quenched with 3% hydrogen peroxide and Background Buster (Innovex Biosciences; Richmond, CA) was used to block non-specific staining. For E-cadherin staining, sections were incubated with rabbit anti-E-cadherin (clone 24E10, Cell Signaling Technology; Danvers, MA) at 1:400 for two hours and then incubated for 30 minutes with Rabbit Envision HRP System reagent (Agilent, Santa Clara, CA). For vimentin staining, sections were incubated with vimentin primary antibody at 1:2000 for 60 min, followed by biotinylated anti-rabbit antibody and Vectastain ABC reagent (Vector Laboratories, Newark, CA). Slides were developed with DAKO DAB plus for five minutes followed by DAB Enhancer (Agilent, Santa Clara, CA) for three minutes before counterstaining with hematoxylin.

## Results

### E-cadherin expression levels do not determine metastatic capacity in the 4T1 stage IV breast carcinoma model

Multiple studies have reported the seemingly paradoxical expression of E-cadherin in aggressive/metastatic 4T1 breast carcinoma cells [17,25,30,31,33–35]. Here, we directly evaluated the role of E-cadherin in either promoting or restraining metastasis in this model. To determine the relative levels of E-cadherin, we immunostained 4T1 cells in comparison with the less metastatic sublines (4T07, 168FARN, and 67NR). These lines were co-isolated with 4T1 from a spontaneous tumor in a Balb/C mouse [36]. The 4T1 line featured prominent E-cadherin expression, while E-cadherin expression was reduced in the next most metastatic 4T07 cell line (Fig. 1A and B). The two least metastatic cell lines, 168FARN and 67NR, had no detectable E-cadherin expression (Fig. 1C and D). Cell surface labeling followed by E-cadherin immunoprecipitation and western analysis confirmed the immunostaining results (Fig. 1E). Thus, paradoxically, higher levels of E-cadherin expression *in vitro* correlate with increasing metastatic potential in this cell set, with the highest expression in the most metastatic 4T1 line.

**Fig. 1.**
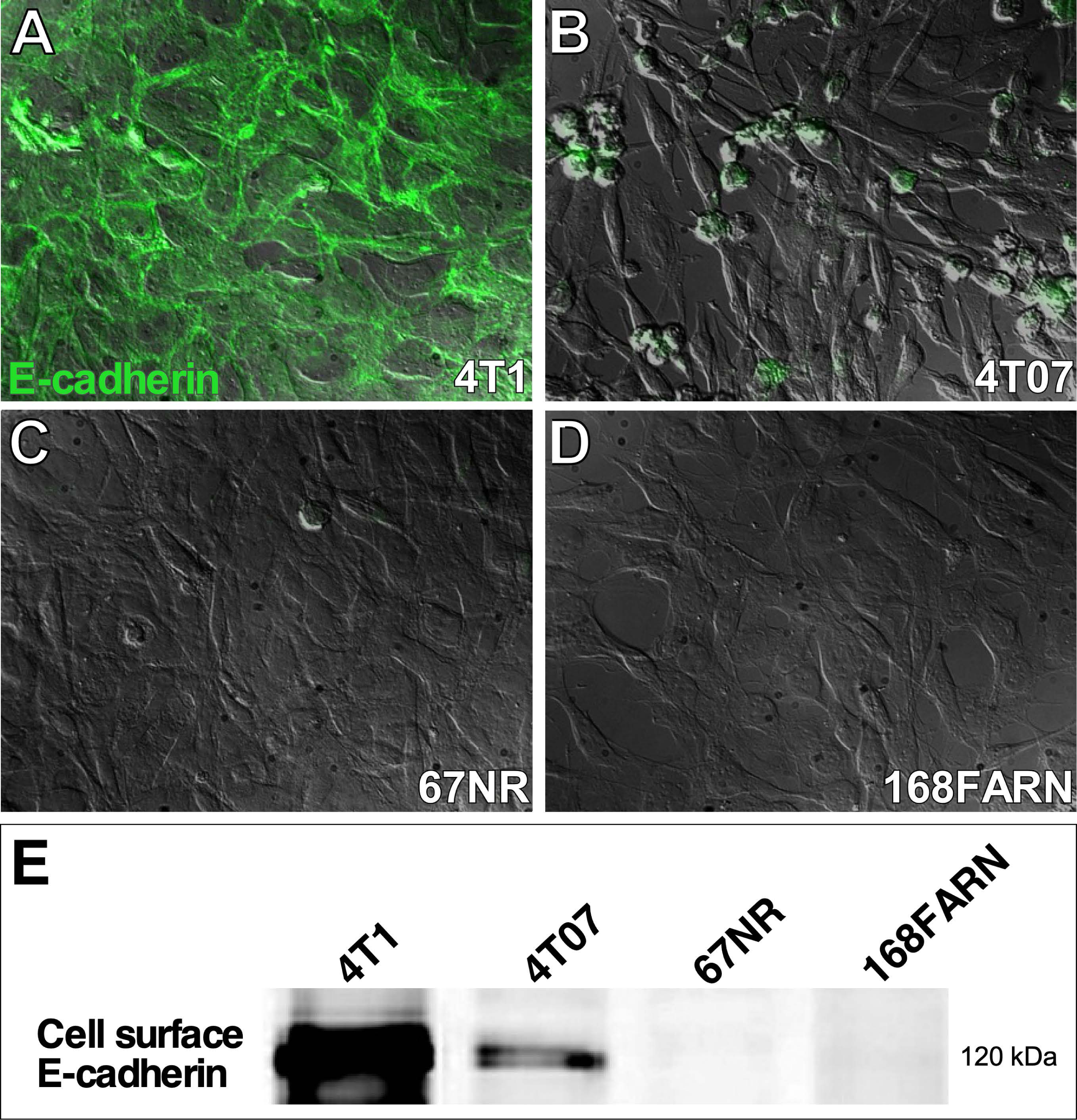
E-cadherin expression correlates with increasing metastatic potential in 4T1 mouse mammary carcinoma cells and three co-isolated sublines. E-cadherin expression was examined by immunostaining in (**A**) 4T1, (**B**) 4T07, (**C**) 67NR, and (**D**) 168FARN cells. Total E-cadherin staining (green) of permeabilized cells is shown superimposed over DIC images. (**E**) Cell-surface biotinylated cells were lysed with Brij 96V, and E-cadherin was immunoprecipitated, followed by western analysis and detection of cell surface-labeled E-cadherin with NeutrAvidin using a LI-COR near-infrared gel imager.

It is possible that E-cadherin has the potential to suppress metastasis of primary tumor cells and promote outgrowth of disseminated cells. To directly test the role of E-cadherin in regulating metastasis we created 4T1 cells with either stable shRNA knockdown of E-cadherin or constitutive expression of E-cadherin possessing a C-terminal myc epitope tag Prior studies have shown that the addition of a myc tag to the C-terminus of E-cadherin does not interfere with either E-cadherin localization or interactions with cytoplasmic protein partners such as catenin [37]. By generating cells with reduced E-cadherin and an enforced expression of E-cadherin we at least partially uncoupled expression levels from transcriptional regulation by mesenchymal transcription factors.

The E-cadherin knockdown cells (Ecad-KD) showed an ∼80% reduction in the level of E-cadherin compared to controls (Fig. 2A, lane 2). Interestingly, the level of β-catenin also was reduced in the Ecad-KD cells compared to parental or Ecad-myc cells (Fig. 2A, lane 2). In some settings, the loss of E-cadherin may free the pool of E-cadherin-associated β-catenin to translocate to the nucleus and function as an EMT-promoting transcriptional coactivator [38,39], although this remains a topic of debate [40]. However, we did not observe an obvious accumulation of nuclear β-catenin in the E-cadherin knockdown cells (Fig. S1). Collectively, these results suggest that, in the 4T1 model, the β-catenin protein liberated by the loss of E-cadherin is mostly degraded, rather than being stabilized as a nuclear transcription factor, at least under standard cell culture conditions.

**Fig. 2.**
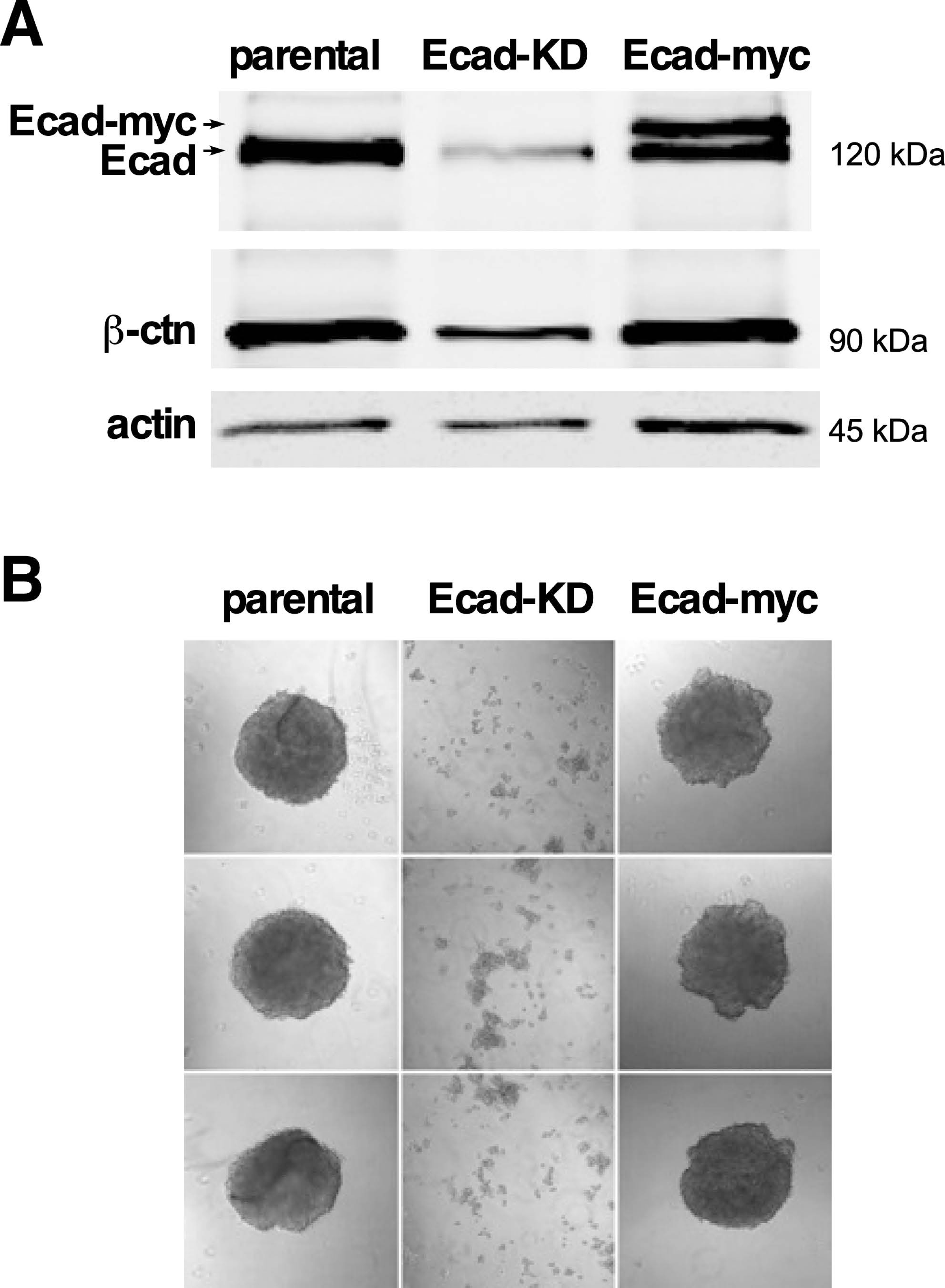
Depletion of E-cadherin in 4T1 cells leads to a reduction in β-catenin and loss of E-cadherin function. (A) Equivalent amounts of NP40 lysates of 4T1 cells (parental), Ecad-KD, and Ecad-myc cells were loaded onto a SDS page gell. The proteins were size separated, transferred to membrane, and incubated with antibodies to the proteins indicated. The primary antibodies were recognized by fluorescently labeled secondary antibodies and the membrane was imaged using the Li-Cor system (company and location). The upper band in lane 3 corresponds to the anticipated size of myc-tagged E-cadherin. **(B)** Parental, Ecad-KD, and Ecad-myc cell spheroids were recovered by pipetting and plated in 35 mm dishes (see Materials & Methods) for immediate photography. Note that Ecad-KD resulted in the loss of spheroid integrity. Knockdown of E-cadherin severely compromised the spheroid stability, indicating strong downregulation of cell-cell adhesion.

In the constitutive expressing (Ecad-myc) cells, surprisingly, the total level of E-cadherin remained nearly the same as that in the parental cells. However, approximately half of the endogenous E-cadherin appeared to be replaced with myc-tagged E-cadherin expressed from the constitutive retroviral promoter (Fig. 2A, lane 3). A possible explanation for the inability of our constitutive expression strategy to increase the total level of E-cadherin expression in 4T1 cells might be that exogenous and endogenous E-cadherin compete for a limited pool of β-catenin, p120-catenin, or vinculin, each of which is required for optimal E-cadherin cell surface localization [41–43]. Thus, endogenous E-cadherin cytoplasmic partners might place limits on the total amount of E-cadherin that can be expressed in 4T1 cells. Regardless, the Ecad-myc 4T1 cells express a substantial proportion of E-cadherin under the control of a constitutive promoter that should no longer be susceptible to transcriptional downregulation via mesenchymal transcription factors such as ZEB1, whereas Ecad-KD 4T1 cells have lost a significant proportion of their E-cadherin expression prior to any induced EMT-like event. Thus, these cell lines were suitable for a functional test of the role of E-cadherin in 4T1 cell metastatic progression.

As a preliminary assessment of E-cadherin function in our cell lines, we performed spheroid forming assays by culturing cells in wells coated with a non-adherent poly-HEMA substrate. Upon recovery by gentle pipetting, the parental and Ecad-myc cells both yielded shear-resistant spheroids, while the Ecad-KD cells could only be recovered as a mixture of single cells and small clumps (Fig. 2B). The loss of spheroid-forming capacity revealed a major functional impact of reducing the level of E-cadherin levels in the Ecad-KD cells.

To directly test the extent to which E-cadherin expression influences metastasis in the 4T1 model, we orthotopically implanted parental, Ecad-KD, Ecad-myc, and vector control cells in Balb/C mice and monitored tumor growth by caliper measurements. The four cell lines displayed similar growth rates at the primary site (Fig. 3A), and *ex vivo* bioluminescence imaging of lungs at the assay endpoint revealed no obvious changes in metastatic capacity upon either a reduction or constitutive expression of E-cadherin (Fig. 3B). We considered the possibility that the similar lung metastatic outgrowth of Ecad-KD cells might reflect re-expression of E-cadherin by a population of cells escaping E-cadherin knockdown. However, measurements of E-cadherin levels and its associated partner, β-catenin, in protein extracts from metastatic tumor cells explanted from the lungs confirmed that knock down of E-cadherin and the concomitant loss of associated β-catenin were maintained after *in vivo* passaging and recovery of the E-cad-KD cells (Fig. 3C). Collectively, our data indicated that expression of the E-cadherin-β-catenin complex *in vitro* is not a strong determinant of 4T1 cell metastatic capacity *in vivo*.

**Fig. 3.**
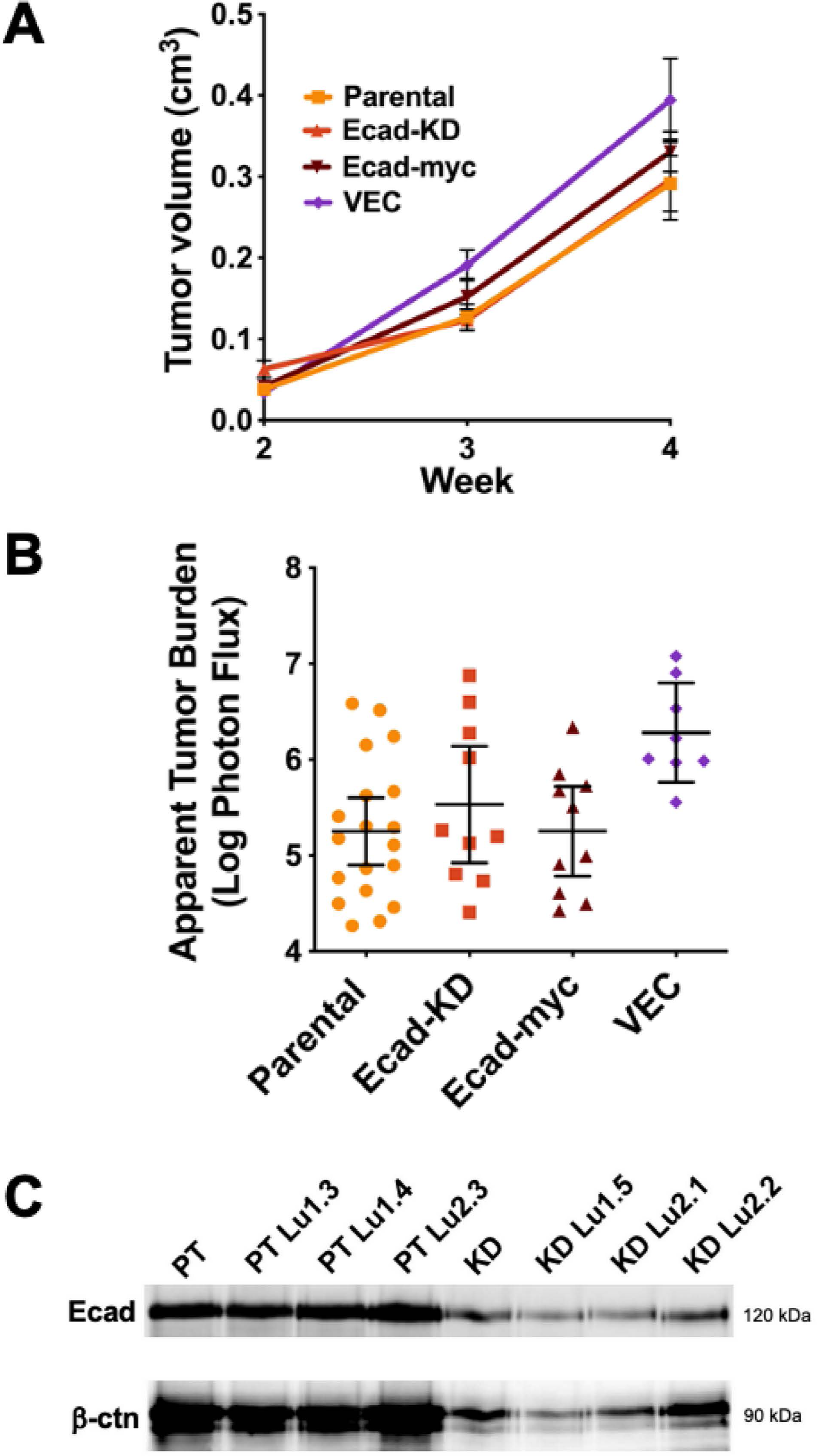
The expression level of E-cadherin-β-catenin complex does not determine the metastatic capacity in the 4T1 breast carcinoma model. (A & B) The pooled results of two separate experiments, in which parental and Ecad-KD, or parental, Ecad-myc, and VEC cells (5 x 10^3^) were orthotopically implanted in Balb/C mice. **(A)** Tumor growth at primary injection sites was monitored by caliper measurements over a four-week period and the tumor volume was plotted. **(B)** The average apparent lung tumor burden for each group was measured at the assay endpoint, as described in Materials and Methods. **(C)** Cells from lung metastatic tumors of parental (PT) and Ecad-KD cells (KD) implanted in mice were explanted as described in Materials and Methods, and cell-surface labeled with biotin. E-cadherin levels were measured by immunoprecipitation, followed by immunoblotting and detection with NeutrAvidin DyLight 800 (top panel). Co-precipitating β-catenin was detected with mouse anti-β-catenin, followed by goat anti-mouse Alexa Fluor 680 (bottom panel). Imaging was by LI-COR near-infrared gel imager. Three lung tumor explanted cell lines from both cell types are shown.

### E-cadherin-based cell-cell junctions are perturbed in 4T1 cells

The fact that *in vitro* E-cadherin expression levels do not influence 4T1 metastatic capacity *in vivo* suggested that E-cadherin function might be altered in these cells. To examine this possibility, we compared the organization of E-cadherin-based cell-cell junctions in 4T1 cells to that of the poorly metastatic, E-cadherin-positive A431 carcinoma cells. In contrast to A431 cells, which display well-organized, circumferential cortical actin, much of the 4T1 cell actin cytoskeleton is in the form of actin cables, aligned perpendicular to cell-cell contacts (Fig. 4). Moreover, 4T1 cell E-cadherin-based junctions are jagged and irregular, compared to the orderly A431 cell junctions (Fig. 4; see also Figure S1). In conjunction with their poor organization, 4T1 cell junctions also display highly dynamic remodeling. Time-lapse video-microscopy revealed that 4T1 cells constantly break and reform cell-cell contacts, even over relatively short observation periods (Fig. 5 and Video S1), as also reported by Elisha et al [31]. Thus, although 4T1 cells can form E-cadherin-dependent, shear-resistant spheroids (Fig. 2B), they are able to rapidly remodel their cell-cell contacts in 2D cultures.

**Fig. 4.**
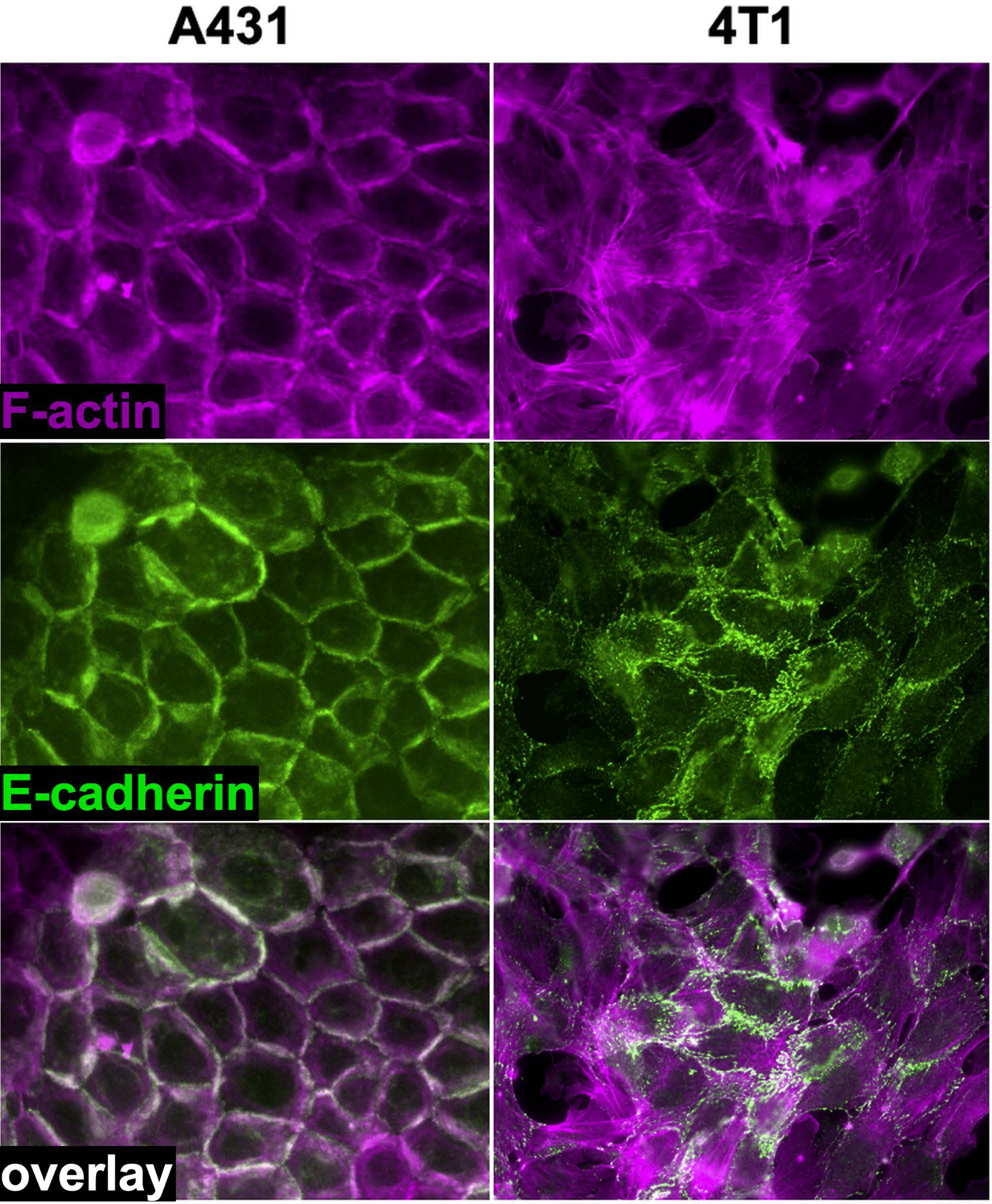
E-cadherin-based cell-cell junctions are disorganized in 4T1 cells compared to those of the poorly metastatic A431 cell line. A431 and 4T1 cells were stained for actin (AlexaFluor 594-conjugated phalloidin; magenta) and E-cadherin (green fluorescence).

**Fig. 5.**
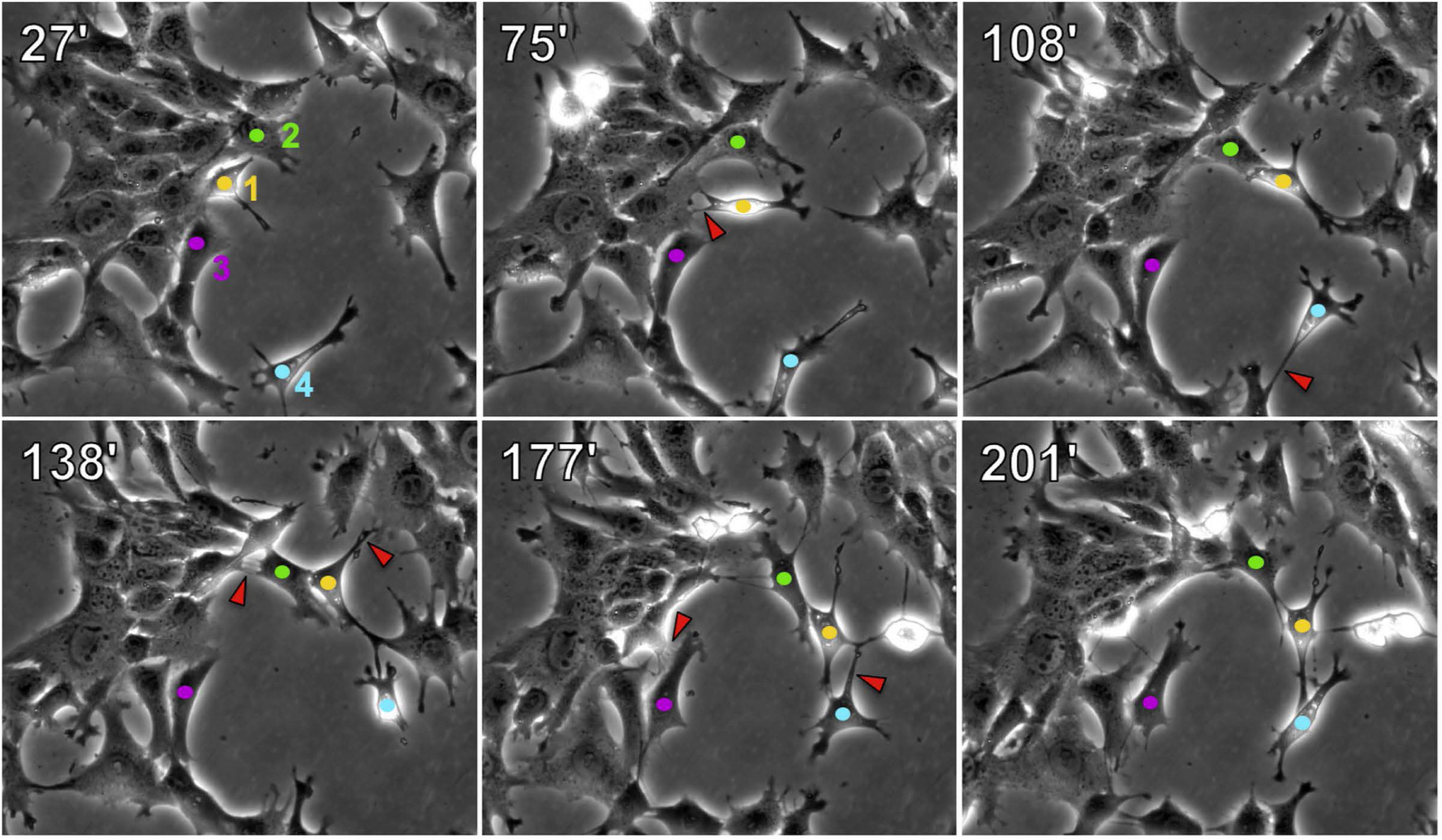
4T1 cell-cell contacts are highly dynamic in 2D cultures. 4T1 cells growing in 35 mm dishes were monitored by time-lapse microscopy for 6 hours. Time of frames in minutes post start of the video are listed in the upper left corner of each panel. Four individual cells are marked in each frame with colored dots (1 = yellow, 2 = green, 3 = magenta, 4 = teal). Red arrowheads mark cell-cell contacts that have separated in the subsequent time frame.

To examine the stability of 4T1 cell-cell junctions in 3D cultures, we created spheroids as in Fig. 2, and embedded them in a 3D collagen matrix. After overnight culture, both parental and Ecad-myc cell spheroids were surrounded by a halo of single cells and small groups of cells that had invaded into the collagen matrix (Fig. 6A,B). By one week post embedding, both types of spheroids displayed extensive invasion, with single cells and multicellular structures emanating from the spheroid (Fig. 6A,B). Ecad-KD cells were not used in this assay as intact Ecad-KD cell spheroids could not be recovered (Fig. 2B).

**Fig. 6.**
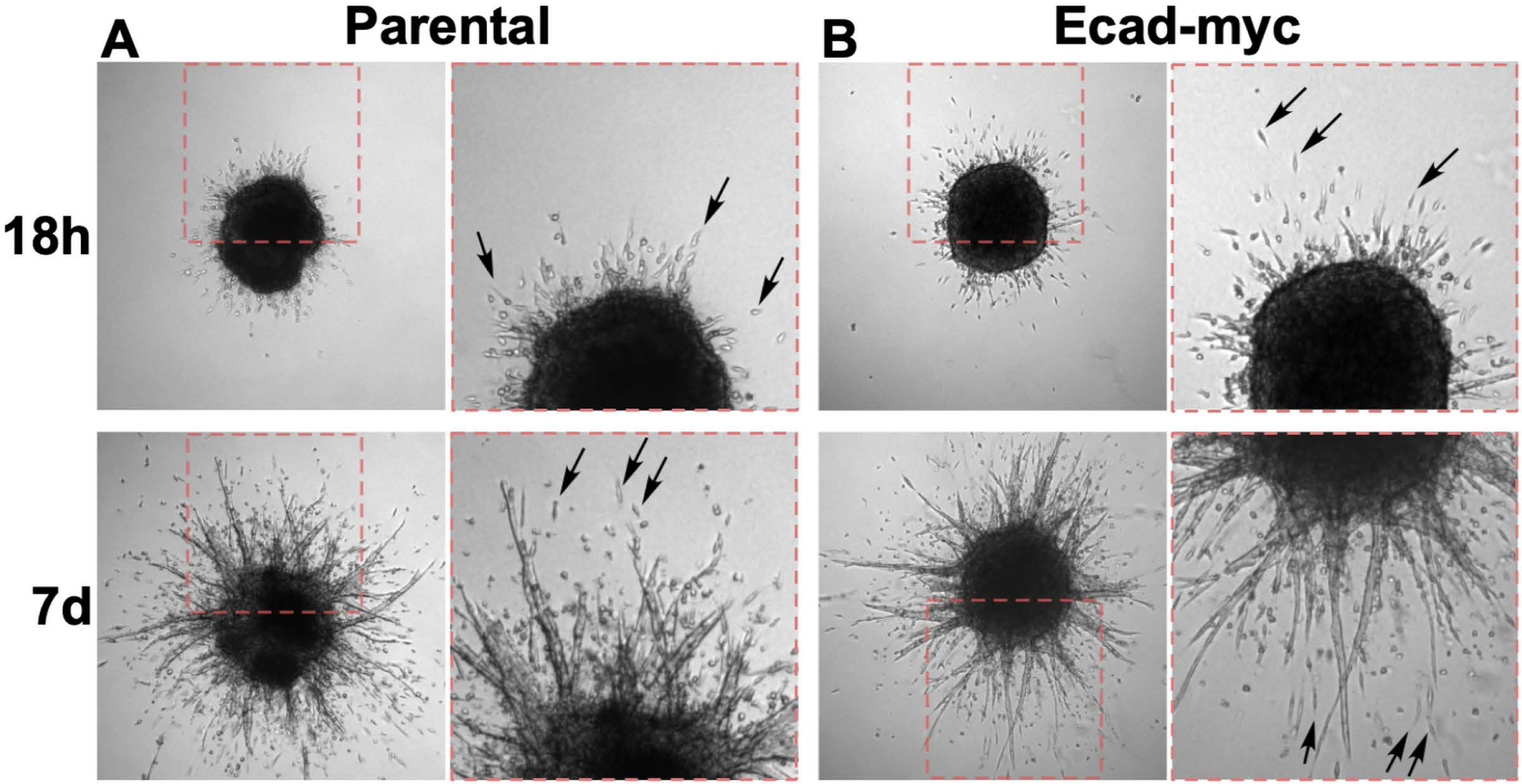
Parental and Ecad-myc 4T1 cell spheroids display extensive invasion of single cells and multicellular structures in 3D collagen. Spheroids of parental (A) and Ecad-myc (B) 4T1 cells were prepared as described in Materials & Methods. After embedding in 3D collagen, micrographs of full spheroids were taken at 18 hours and 7 days. Dashed red boxes in left hand panels indicate the regions magnified in the adjacent panels. Arrows point to examples of single cells that have detached from the spheroids during invasion.

The ability of individual 4T1 cells to break free from a spheroid and invade a 3D collagen matrix within 18 hours could be explained by an EMT-like event, in which the most invasive cells downregulated E-cadherin. Therefore, we investigated the timing of the onset of invasion of parental 4T1 cells by time-lapse video-microscopy. As early as two hours after embedding, invasive cells oriented towards the collagen matrix were evident at the spheroid boundary (Fig. 7). By as early as five hours after embedding, the first singly invading cells had fully emerged from the spheroid (Fig. 7; see also Video S2). The process by which single cells and small groups of cells emerge from the spheroid can be monitored in detail through video recordings (Video S3, a zoomed in field of Video S2). Invading cells appear to overcome significant traction forces exerted by the spheroid, which often appears to pull emerging cells back into the spheroid. Eventually, individual cells or small clusters of cells free themselves from the spheroid, often emerging from the tips of strands of collectively invading cells (Video S3), like the invasive structures described by Cheung et al. in tumors and organoids from the MMTV-PyMT breast cancer model, as well as primary human breast tumor explants in 3D collagen I [44]. To evaluate E-cadherin expression in the invading cells, we fixed and immunostained E-cadherin in the spheroid used to create Figure 7 and Videos S2 and S3. E-cadherin expression was evident in both single cells and small groups of cells invading in contact with each other (Fig. 8). Negative control staining of another spheroid showed only a low background of non-specific signal (Fig. 8). Thus, a loss of E-cadherin was either extremely transient or unnecessary for cell escape and invasion in this model.

**Fig. 7.**
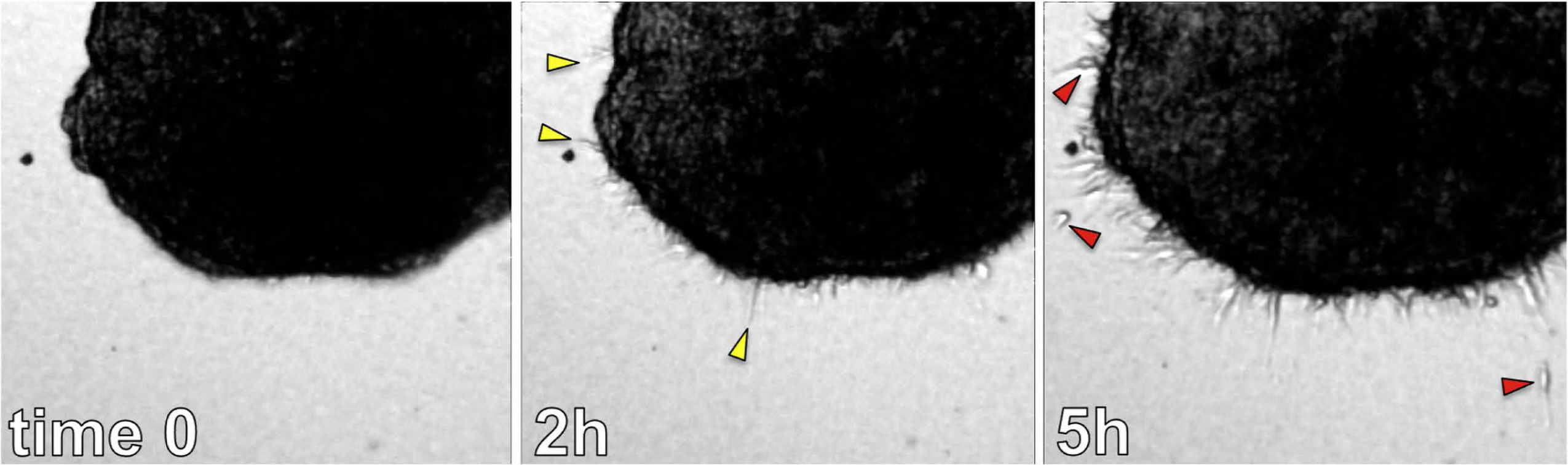
4T1 spheroids display early onset single cell invasion in 3D collagen. 4T1 spheroids were prepared as described in Materials & Methods. Micrographs of the edge of a spheroid are shown in which the image was taken immediately after embedding, and at two and five hours after embedding. The yellow arrowheads in point to multicellular structures oriented toward the matrix at the 2-hour time point. Red arrowheads in point to single cells that have emerged from the spheroid by the 5-hour time point. This sequence illustrates how some individual cells can rapidly dissociate from the spheroid within hours after embedding in a collagen matrix.

**Fig. 8.**
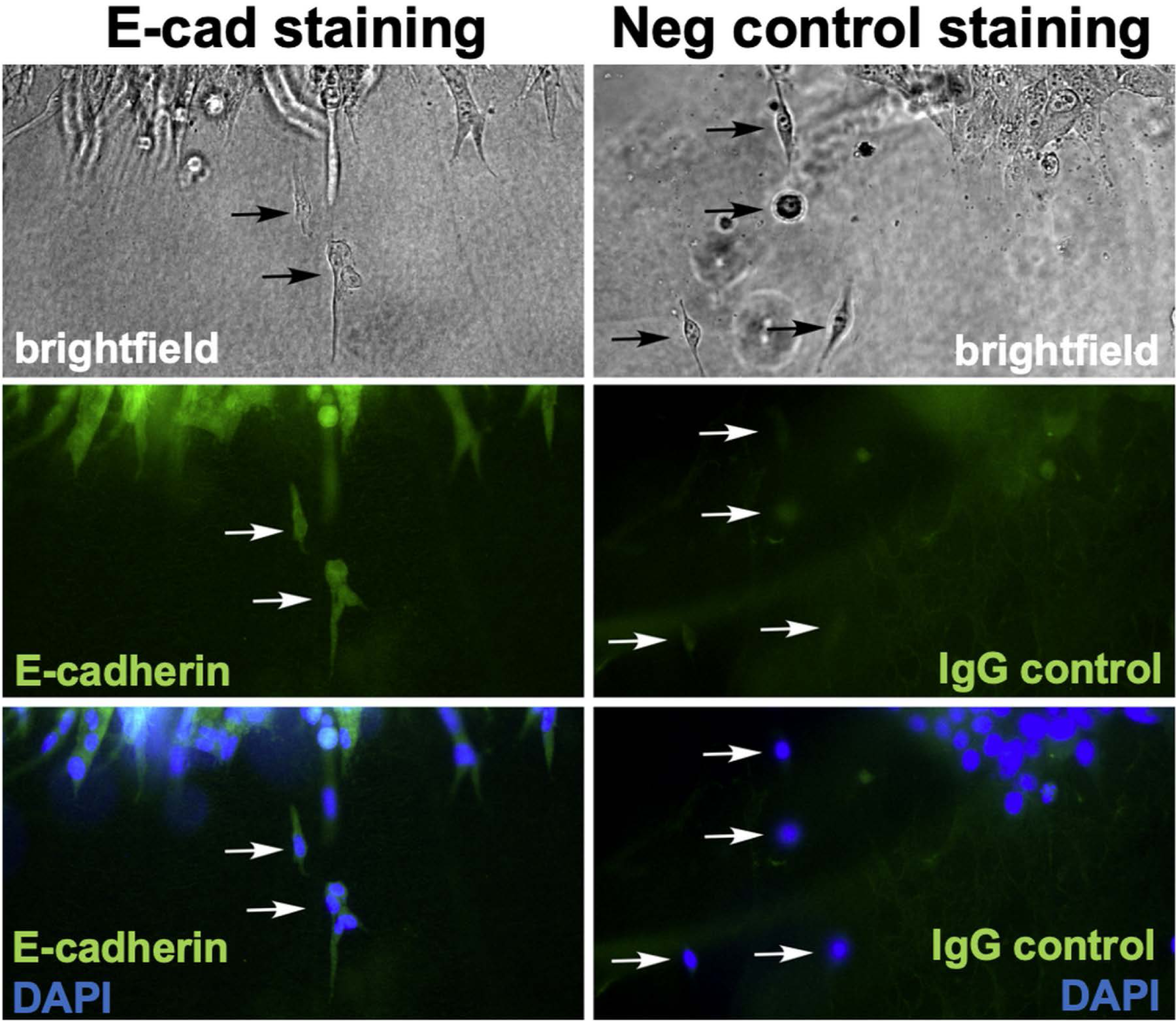
E-cadherin is retained in single cells and clusters of cells emerging from 4T1 cell spheroids embedded in 3D collagen. The spheroid depicted in Figure 7 and Videos S2 and S3 was fixed after the 40-hour time-lapse experiment, and stained with an anti-E-cadherin antibody. Another spheroid from a parallel experiment was fixed and stained with a mouse IgG isotype negative control antibody. For both samples, cell nuclei were visualized with DAPI, and the edges of the spheroids were imaged by brightfield and fluorescence microscopy. Arrows point to single cells or small groups cells that have emerged from the spheroids. These results demonstrate that cells and cell clusters invading into the 3D collagen matrix either retained E-cadherin expression throughout the invasion process, or they rapidly re-expressed E-cadherin upon dissociation from the spheroid.

Collectively, these observations suggest that E-cadherin’s ability to enforce key features of an epithelial phenotype is disrupted in 4T1 cells, facilitating their invasion without loss of E-cadherin.

### 4T1 cells display mixed expression of p120-catenin isoforms 1 & 3 consistent with a hybrid E/M phenotype

A potential explanation for the inability of E-cadherin to suppress 4T1 cell invasion is a lack of E-cadherin interaction with cytoplasmic partners, α-catenin, β-catenin, and p120-catenin. However, co-immunoprecipitation experiments confirmed that α-catenin and β-catenin are expressed in 4T1 cells and co-immunoprecipitate with E-cadherin (Fig. 9A,B; see also Fig. 3C, which showed β-catenin/E-cadherin co-IPs). The p120-catenin is also expressed in 4T1 cells and co-immunoprecipitates with E-cadherin. Interestingly, for p120-catenin, the mesenchymal isoform 1 and epithelial isoform 3 are co-expressed in 4T1 cells (Fig. 9C). Isoform 3 of p120-catenin predominates in epithelial cells, and induction of EMT, for example by forced Snail expression, can trigger a switch from isoform 3 to isoform 1 [45,46]. Thus, the presence of both isoforms suggests a hybrid epithelial-mesenchymal (E/M) phenotype. Support for this idea comes from several additional studies, which collectively paint a picture of a hybrid epithelial-mesenchymal phenotype for the 4T1 cell line (see Discussion and Supplemental Table 2).

**Fig. 9.**
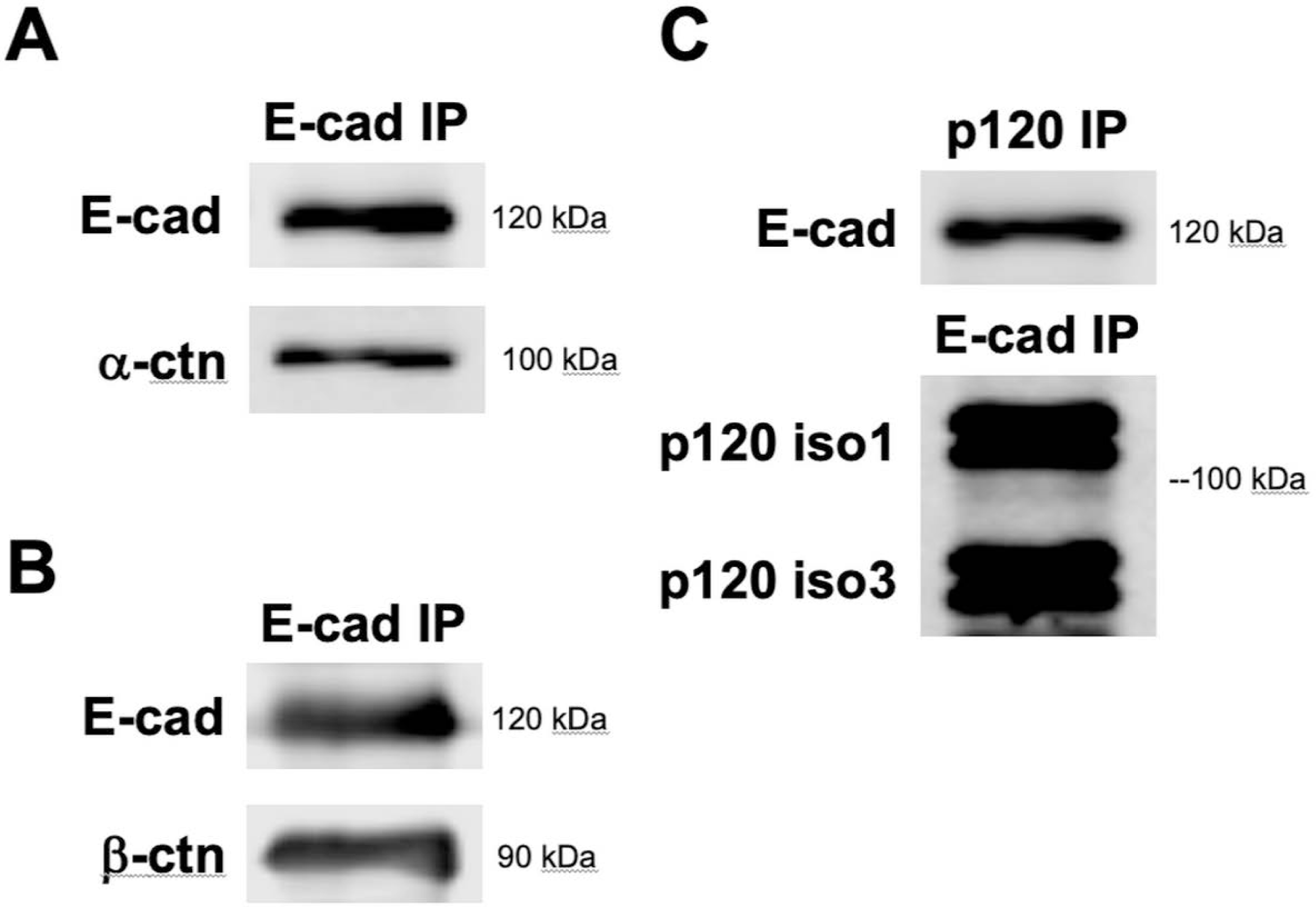
E-cadherin co-immunoprecipitates with its cytoplasmic partners α-catenin, β-catenin, and p120 catenin isoforms 1 and 3 in 4T1 cells. NP40 lysates of 4T1 cells were immunoprecipitated for either E-cadherin (**A, B, & C**) or p120 catenin (**C**). The precipitated proteins were used in western analysis with the primary antibodies as shown, followed by detection with fluorescently labeled secondary antibodies and LI-COR near-infrared gel imaging.

### A hybrid E/M phenotype and EMT-like changes in sub-regions of a tumor may contribute to 4T1 cell metastasis *in vivo*

To further investigate the hybrid E/M phenotype of 4T1 cells *in vivo*, we immunostained for E-cadherin in primary tumors and lung metastatic cells *in vivo*. E-cadherin expression initially appeared widespread throughout the primary tumor tissue (Fig. 10A). However, upon higher magnification, it was apparent that there were areas in which a subset of tumor cells appeared E-cadherin-negative, even when juxtaposed to areas of E-cadherin positive cells (Fig. 10B). We also analyzed 4T1 tumors by immunostaining for the mesenchymal marker, vimentin. (Fig. 10C). Unlike E-cadherin, which was variably expressed, vimentin expression appeared uniform throughout 4T1 primary tumors, as previously reported [47]. By constrast, vimentin staining was absent from an internal negative control of mammary epithelial cells captured in the same tissue slice (Fig. 10C). Thus, primary 4T1 cell tumors contain populations of cells that express both E-cadherin and vimentin as well as populations that lack E-cadherin and express vimentin.

**Fig. 10.**
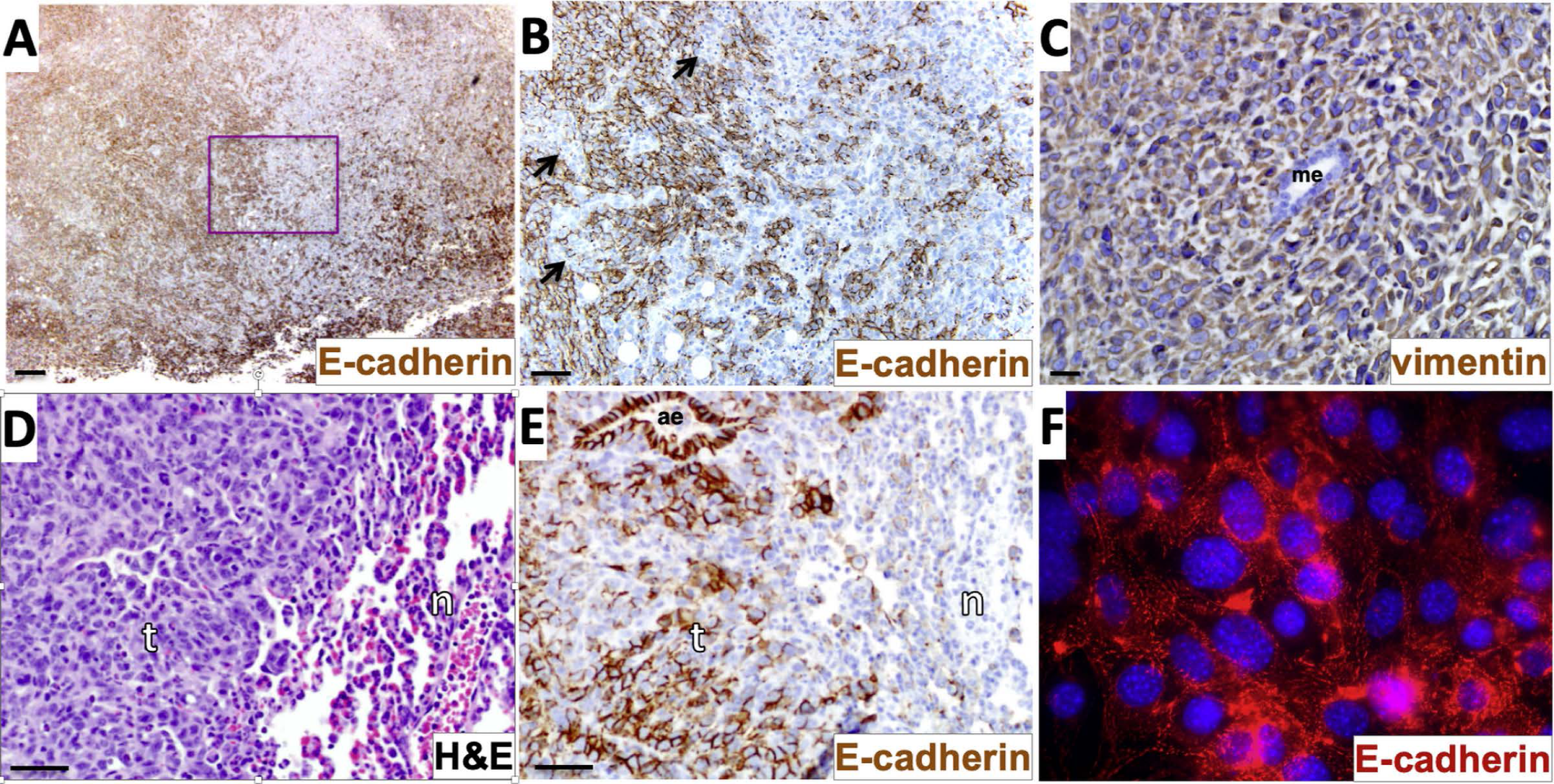
4T1 primary tumors and lung metastatic tumors display both E-cadherin positive and negative regions, consistent with EMT and a hybrid E/M phenotype. (**A & B**) Primary tumor tissue was stained with E-cadherin immunohistochemistry (brown) and counterstained with hematoxylin. The area outlined by magenta in panel **A** is enlarged in panel **B**. Arrows indicate E-cadherin-negative zones within an overall E-cadherin-positive sector of the tumor (**B**). (**C**) Primary tumor tissue was stained with vimentin immunohistochemistry (brown) and counterstained with hematoxylin. Tissue from lung metastatic tumors were stained with H&E (**D**) or with E-cadherin immunohistochemistry and hematoxylin counterstain (**E**). (**F**) Cells explanted from a 4T1 metastatic lung tumor as described in Materials & Methods were immunostained with anti-E-cadherin (red) and DAPI (blue). Labels in (**C, D, & E**) are me, mammary epithelium; t, tumor tissue; n, normal lung tissue; ae, airway epithelium. Scale bars in (**A**) = 200 μm, in (**B, D, & E**) = 50 μm, and in (**C**) = 20 μm.

To determine if this heterogeneity in E-cadherin expression also exists in metastases, lung metastatic tumor tissue (Fig. 10D) was immunostained with an antibody to E-cadherin. E-cadherin expression was variable within lung-metastatic tumor tissue, but clearly present within the nearby internal positive control airway epithelial cells within the same tissue slice (Fig. 10E). Thus, as we observed in primary tumors, both E-cadherin positive and negative cells were found in metastatic tumors. By contrast, when lung metastatic cells were recovered and cultured *in vitro* (selecting for tumor cells based on G418 resistance, a feature of the *luciferase* expression vector), universal E-cadherin expression was observed by the time a bulk population of explanted tumor cells was recovered (Fig. 10F). This suggests that retention or re-expression of E-cadherin is a feature of cells that can grow out more efficiently in culture following explantation. Taken together, these data support a scenario in which E-cadherin is downregulated in a subset of cells in 4T1 primary tumors, which are already composed of cells that possess a hybrid E/M phenotype that could facilitate collective invasion and dissemination without E-cadherin downregulation. This combination of altered cell adhesion proteins and hybrid cellular identity is a feature consistent with the aggressive metastatic behavior of the 4T1 carcinoma model.

## Discussion

E-cadherin has classically been viewed as a metastasis suppressor due to its ability to enforce a sessile, epithelial phenotype with strong, organized cell-cell junctions and attenuation of β-catenin signaling [23–26,38–40]. Triggering an EMT can result in downregulation of E-cadherin that promotes tumor cell invasion and metastasis by releasing the anti-migratory constraints of cell-cell contact. Conversely, at least in some settings, MET may be required at the metastatic site to facilitate macroscopic tumor outgrowth [14–21]. E-cadherin expression is often used as a marker to monitor EMT and MET events; however, the extent to which E-cadherin itself is required to restrain invasion at the primary tumor site or to promote growth upon MET at a secondary tumor site remains the subject of ongoing investigation. Here we found that neither depletion nor constitutive expression of E-cadherin had an obvious impact on 4T1 tumor growth or metastasis *in vivo*, adding to the evidence that acquiring metastatic and non-metastatic properties are complex processes that rely on more than E-cadherin as a switch.

Our results contrast with those of Chu et al., who reported that RNAi silencing of E-cadherin in the SUM149 and Mary-X inflammatory breast cancer cell lines abrogated growth *in vivo* [30]. Depleting E-cadherin by RNAi in 4T1 cells was also reported to block primary tumor growth in this study. One possible explanation for this discrepancy is that our 4T1 E-cadherin-knockdown cells were maintained as a polyclonal population, while Chu et al. isolated a subclone, which potentially might have been more strongly dependent on E-cadherin for growth than the bulk 4T1 cell population.

Our results also contrast somewhat with those of Elisha et al., who reported a modest reduction in 4T1 cell lung metastatic growth upon E-cadherin knockdown, when observed at early time points (∼1-2 weeks after tail vein or orthotopic injection) [31]. In our study, both E-cadherin-depleted and E-cadherin-myc expressing cells displayed a similar growth rate compared to control cells at the primary site and no obvious difference in lung colonization at a later timepoint (after ∼4 weeks). Our observations agree with a report, in which complete deletion of E-cadherin via CRISPR gene editing had no obvious impact on 4T1 cell metastatic capacity [48]. However, our results also do agree with Elisha et al. with regards to the dynamic nature of 4T1 cell-cell junctions and the possibility that modes of collective cell migration involving E-cadherin may contribute to 4T1 cell metastasis [31]. On balance, the available data suggest that E-cadherin expression in the bulk 4T1 cell population has only a modest direct impact on the metastatic capacity of the cells, at least when measured at later time points, rendering the cells largely E-cadherin-indifferent regarding the ultimate outcome of disease progression in this model. Our ability to deplete E-cadherin by ∼80% without blocking macroscopic outgrowth may reflect either that the ∼20% residual E-cadherin expression in the E-cadherin-depleted cells was sufficient to support outgrowth or that expression of E-cadherin by itself is not the critical aspect of the epithelial phenotype that serves to promote macroscopic outgrowth.

Although 4T1 cells retain E-cadherin expression *in vitro* and form E-cadherin-dependent, shear-resistant spheroids, immunostaining and time-lapse video microscopy revealed that 4T1 cell-cell junctions are disorganized and highly dynamic (Figs. 4,5 and Videos S1-S3). The ability of E-cadherin-positive 4T1 cells to dissociate from tumor spheroids is reminiscent of the mechanical disruption of cell-cell junctions upon treatment of Madin-Darby canine kidney epithelial (MDCK) cells with hepatocyte growth factor (HGF) [49]. In this study, HGF triggered MDCK cell scattering without downregulation of E-cadherin. Instead, HGF activated a contractile actin cytoskeleton, with actin bundles terminating orthogonally at cell-cell boundaries, like what we observed in 4T1 cells. The increased actomyosin contractility in response to HGF caused MDCK cell-cell junctions to be pulled apart, with no detectable reduction in cell surface E-cadherin expression. We propose that 4T1 cell junctions are constitutively disorganized and dynamic, contributing to their invasive and metastatic phenotype despite their retention of E-cadherin expression.

The 4T1 murine breast carcinoma is a widely utilized model of metastatic breast cancer because it is one of few cell lines that features rapid, robust spontaneous metastasis from an orthotopic site in an immunocompetent host. Prior studies have reported 4T1 EMT events *in vivo*, and the isolation of circulating and disseminated 4T1 cell sub-lines with both more purely mesenchymal and hybrid-E/M molecular phenotypes [35,47]. The functional significance of 4T1 cells possessing a hybrid epithelial and mesenchymal phenotype is supported by numerous studies, which at the time they were performed, were not directly aimed at establishing an E/M phenotype in these cells (Supplementary Table 2). A landmark study describing the expression of the EMT-inducing transcription factor Twist in 4T1 cells helped to solidify the concept of EMT as a critical determinant of metastasis [29]. In addition to Twist, EMT-promoting transcription factor Zeppo1 may also contribute to 4T1 cell metastasis [50]. However, we and others noted retention of E-cadherin and epithelial-specific splicing factor ESRP1 in the 4T1 model [17,30,31,48,51]. Analyses of microRNA expression in the 4T1 cell system have also revealed co-expression of both pro-epithelial miRNAs (miR-200 family and miR-155) and pro-mesenchymal miRNAs (miR-9, miR10b) [17,34,52–54]. The co-expression of epithelial and mesenchymal master regulators may be critical for the aggressive phenotype of 4T1 cells since depleting ESRP1, Twist, Zeppo1, miR-9, or miR-10b all suppress 4T1 metastasis [29,50–53], while over-expressing miR-200 family members promotes metastasis of the closely related 4TO7 breast carcinoma cells, which normally fail to form macroscopic colonies in lung [17,34]. Moreover, altering expression of “phenotypic stability factors” OVOL2 or GRHL2, both of which may help to enforce a stable E/M phenotype [55], also reduced metastatic capacity in the 4T1 model [56,57]. Lastly, Snail has been proposed as an EMT-TF that promotes a hybrid E/M state and increased metastatic capacity [58], and manipulating Snail protein expression up or down by forced expression or RNAi-knockdown of ubiquitin E3 ligase FBXO11, which targets Snail, provided evidence that increased Snail expression promotes 4T1 cell metastasis [59]. Collectively, these numerous, independent studies strongly support the view that the 4T1 model can manifest a hybrid E/M state of metastatic breast cancer.

An interesting aspect of the hybrid phenotype of 4T1 cells is their co-expression of p120 catenin isoforms 1 and 3. Intriguingly, Zeppo1, which promotes 4T1 cell metastasis, also promoted a shift in p120 catenin expression from the epithelial isoform 3 to the mesenchymal isoform 1 in non-tumorigenic mammary epithelial cells cultured in 3D [50]. Inducers of a mesenchymal phenotype can promote dramatic morphological changes even in the ongoing presence of E-cadherin. For example, TGF-β-treated NMuMG mammary epithelial cells undergo a profound morphological transition with a loss of organized cell junctions that precedes the loss of E-cadherin cell surface expression by at least 2 days [60]. In this study, from Wheelock and colleagues, molecular changes in response to TGF-β treatment included disruption of the normal cortical actin cytoskeleton and formation of trans-cellular stress fibers, like the arrangement of the actin cytoskeleton in 4T1 cells. These changes occurred early, despite the initial maintenance of E-cadherin, and the continued association of E-cadherin with its partners, α- and β-catenin [60]. We suggest that in a similar manner, a constitutive, partially mesenchymal phenotype overrides the ability of E-cadherin to maintain organized adherens junctions in 4T1 cells.

If 4T1 cells possess a constitutive, partially mesenchymal phenotype that can override the ability of E-cadherin to restrain detachment and invasion, one might posit that no further EMT-like events would be required to facilitate their metastasis. However, analysis of cells *in vivo* suggests additional layers of complexity. We detected the maintenance of E-cadherin positive 4T1 cells *in vivo*, as previously reported [33]; however, close inspection revealed regions of tumor cells with reduced levels of E-cadherin, which has also been reported in the 4T1 model [35]. Staining of E-cadherin in metastatic colonies revealed a similar picture, with mixed E-cadherin positive and negative cells (Figure 10). Overall, our data suggest that localized E-cadherin downregulation, overlaid onto a phenotype that is already partially mesenchymal, might contribute to the aggressive nature of the 4T1 breast carcinoma *in vivo*. As such, the 4T1 model can serve as an experimental platform to explore potential treatment strategies for invasive carcinomas that retain E-cadherin expression. Such strategies may include restoring the ability of E-cadherin to suppress invasion, as suggested by a study utilizing an E-cadherin activating antibody in the 4T1 model [61]. Conversely, a better understanding of how epithelial master regulators such as miR-200 and miR-155 can sometimes function to promote metastatic outgrowth may lead to novel strategies for selectively targeting those pro-metastatic functions associated with an epithelial phenotype.

## Acknowledgements

This research was supported by a Pilot Grant from the Holden Comprehensive Cancer Center. Cell sorting and flow cytometry were performed at the Flow Cytometry Core Facility at the Holden Comprehensive Cancer Center in the University of Iowa Carver College of Medicine.

## Dedication

This study is dedicated to the memory of Dr. Mary Elizabeth Herndon, PhD – a luminous presence in the lab and in life.

