## Supplemental Figures and Tables for "The 4T1 breast carcinoma model possesses hybrid epithelial/mesenchymal traits linked to a highly metastatic phenotype"

### Supplementary Information

**Figure S1**. **Loss of β-catenin in E-cadherin knockdown cells.** Parental and Ecad-KD cells were cultured on cover slips overnight. Cells were fixed, permeabilized and immunostained for (A) E-cadherin or (B) β-catenin antibodies, together with (A) AlexaFluor 594-conjugated phalloidin (to reveal F-actin) or (B) DAPI (to stain DNA). The level of β-catenin appeared much reduced in the E-cad KD cells compared to parental cells, and the remaining β-catenin did not obviously re-localize to the cell nucleus.

**S1 Video. 4T1 cell junctions are highly dynamic.** This video captures 4T1 cells making and breaking cell-cell contacts over a six-hour period, with one frame every three minutes. The play rate is eight frames/sec (1,440 X real time). A still shot from this video corresponds to Fig. 5 in the manuscript.

**S2 Video. 4T1 cells within a spheroid invade in 3D culture conditions.** This video shows 4T1 cell invasion into a 3D collagen matrix over a 40-h period, with one frame every five minutes. The play rate is eight frames/sec (2,400 X real time). A still shot from this video corresponds to Fig. 7 in the manuscript.

**S3 Video. A magnified view of 4T1 cells within a spheroid invading in 3D culture conditions.** This video is a magnified and cropped view of the S2 video centered around the bottom edge of the spheroid. The play rate is 16 frames/sec (4,800 X real time). Arrows in the final frame indicate cells that correspond to the immunostained image in Fig. 8 of the manuscript.

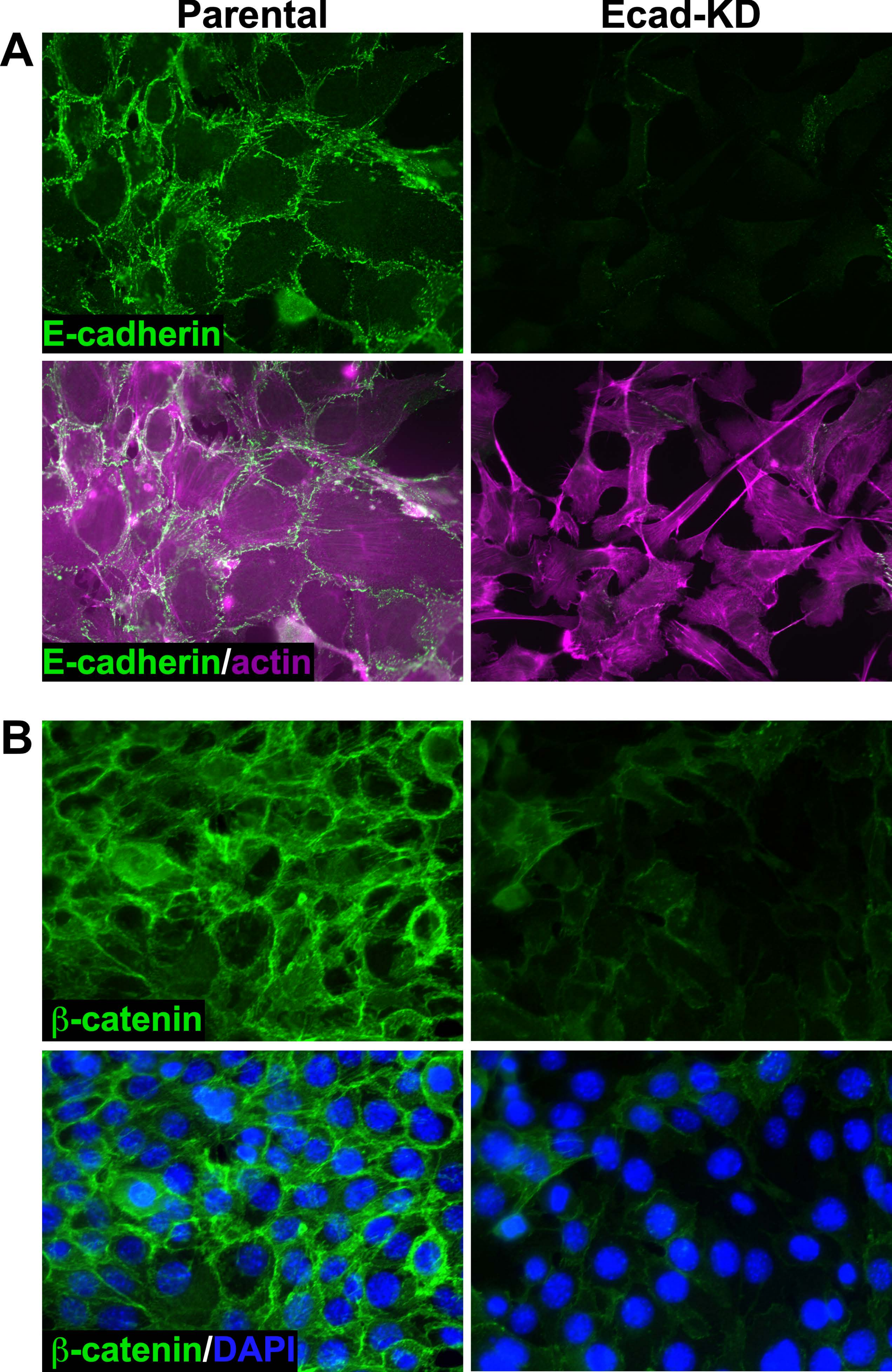
Parental
Ecad-KD
A
E-cadherin
E-cadherin/actin
B
β-catenin
β-catenin/DAPI

**Table S1. Primary antibodies used in this study**

| Target of antibody | Source | Antibody clone/cat# | Type | Reactivity[1] | Application in this study[2] |
|---|---|---|---|---|---|
| E-cadherin | Sigma-Aldrich | DECMA-1 | Rat IgG | H M D | Flow, IF, IP |
| E-cadherin | Thermo Fisher Scientific | Clone 36, BD | Mouse IgG | H M Rt D | IF, IP |
| E-cadherin | Cell Signaling Technology | Clone 24E10 | Rabbit IgG | H M | IHC |
| E-cadherin | Jack Lilien & Janne Balsamo, University of Iowa | Polyclonal Ab to E-cadherin cytoplasmic tail | Rabbit IgG | H M | WB |
| α-catenin | Thermo Fisher Scientific | Clone 5, BD | Mouse IgG | H M Rt | WB |
| β-catenin | Thermo Fisher Scientific | Clone 14, BD | Mouse IgG | H M Rt D | WB, IF |
| p120-catenin | Thermo Fisher Scientific | Clone 98, BD | Mouse IgG | H M Rt D | WB, IP |
| β-actin | Biolegend | Poly6221 | Rabbit IgG | H M Rt | IB |
| Vimentin | Abcam | EPR3776 | Rabbit IgG | H M Rt | IHC |

[1]H, Human; M, Mouse; D, Dog; Rt, Rat

[2]Flow, flow cytometry; IF, immunofluorescence; IP, immunoprecipitation, IHC, immunohistochemistry; WB, western blotting

**Table S2. Studies supporting a hybrid E/M phenotype in the 4T1 Breast Carcinoma Model**

| Epithelial/Mesenchymal Regulator or Marker[1] | Outcome of functional intervention in the 4T1/4T07 breast cancer models |
|---|---|
| Twist (drives mesenchymal phenotype) | Depletion in 4T1 cells *suppressed* spontaneous metastasis [29] |
| miR-9 (targets E-cadherin mRNA) | Downregulation in 4T1 cells *suppressed* spontaneous metastasis [52] |
| miR-10b (upregulated by Twist) | Downregulation in 4T1 cells *suppressed* spontaneous metastasis [53] |
| Zeppo1/ZNF703 (can trigger EMT) | Depletion in 4T1 cells *suppressed* spontaneous metastasis [50] |
| miR-200 family (downregulates Zeb1, enforces epithelial phenotype) | Highly expressed in 4T1 cells. Over-expression in 4T07 cells *enhanced* spontaneous & experimental metastatic colonization [17, 34] |
| ESRP-1/2 (epithelial splicing factors) | Depletion of ESRP-1 in 4T1 cells reduced CD44v expression and *suppressed* spontaneous metastasis [51] |
| E-cadherin | Depletion or forced expression in 4T1 cells had minimal impact on primary tumor growth or spontaneous metastasis (this study)<br>E-cadherin depletion modestly reduced metastatic outgrowth at early time points [31].<br>An E-cadherin-depleted 4T1 subclone showed impaired primary tumor growth [30]<br>Preliminary data indicated that CRISPR deletion of E-cadherin in 4T1 cells had no effect on metastatic growth [48] |
| FBXO11 (promotes degradation of Snail, which may be associated with a hybrid E/M phenotype) | Depletion of FBXO11 increased Snail expression in 4T1 cells and *promoted* 4T1 cell metastasis; over-expression of FBX011 had the converse effect [59] |
| miR-155 (prevents EMT) | Forced expression of miR-155 in 4T1 *suppressed* spontaneous metastasis from fat pad but *promoted* experimental metastasis after tail vein injection [54] |
| OVOL2 (may stabilize hybrid E/M state when expressed at endogenous level) | Forced over-expression increased epithelial phenotype of 4T1 and *suppressed* spontaneous lung metastasis [56] |
| GRHL2 (may stabilize hybrid E/M state when expressed at endogenous level) | Depletion of GRHL2 *suppressed* 4T1 cell lung metastasis [57] |

[1]Mesenchymal markers are shaded **orange**, epithelial markers are shaded **blue**, and phenotypic stability factors are shaded **green**.
